# Simulated climate warming scenarios lead to earlier emergence and increased weight loss but have no effect on overwintering survival in solitary bees

**DOI:** 10.1101/2024.10.18.619057

**Authors:** Jannik S. Möllmann, Liv Lörchner, Dean Hodapp, Ina Knuf, Hongfei Xu, Thomas J. Colgan

## Abstract

Insect pollination is critical for both wildflower stability and agricultural yields, with solitary bees being a group of pollinators of fundamental importance. However, documented declines in populations, exacerbated by environmental pressures, including climate change, pose significant threats to the provision of ecosystem services. Exposure to elevated temperatures during periods of dormancy, such as overwintering, is predicted to lead to phenological shifts, changes in condition, and impacts on survival. However, we currently lack studies that inform how such aspects are affected in future climate change scenarios. Using simulated temperature regimes informed by predictions of the Intergovernmental Panel on Climate Change, we exposed overwintering mason bees (*Osmia* species) to three field-relevant temperature profiles based on either present-day overwintering temperatures or future temperatures predicted under two major climate warming scenarios (SSP2-4.5 and SSP5-8.5) and measured how temperature exposure affected emergence timing, weight loss and survival. We found that exposure to temperatures under intermediate and worst-case climate warming scenarios led to earlier emergence by approximately three and six weeks, respectively, with increasing divergences in timing of emergence between the sexes of *Osmia bicornis*, which may lead to intraspecific phenological mismatches. While we found no effect of temperature on overwintering survival rates, we observed increased weight loss prior to emergence but found that in contrast to other studies, it only mildly mediated shifts in emergence timing brought about by temperature exposure, suggesting that weight loss is unlikely to play a major role as a trigger of emergence timing in mason bees. Our study contributes to the growing literature highlighting the impact that temperatures under climate change models will have on the timing of key life events for essential pollinators, which may have consequences at the population and community levels.

## Introduction

Insect pollination is widely regarded an essential ecosystem service (Kremen et al. 2007), being not only crucial for wildflower stability and biodiversity (Potts et al. 2016; Kremen et al. 2007; Kleijn et al. 2015) but also contributing to agricultural yields (Kremen et al. 2004; Reilly et al. 2020), with economic benefits estimated at over €150 bn (Gallai et al. 2009). Despite these benefits, reported population declines of insect pollinators have raised concerns over the long-term provision of pollination services and the associated implications for both natural ecosystems and food security (Lippert, Feuerbacher, and Narjes 2021; Klein et al. 2007). Several environmental factors have been suggested as key contributors to declines, including habitat loss and fragmentation, increased pesticide exposure, disease, invasive species, as well as climate change (Cameron et al. 2011; Potts et al. 2016; Seibold et al. 2019; Powney et al. 2019; Hallmann et al. 2017). In particular, climatic change has increasingly been highlighted as negatively impacting pollinators (Walther et al. 2002; Vanbergen and the Insect Pollinators Initiative 2013; Scaven and Rafferty 2013; Harvey et al. 2023) through heightened exposure to elevated frequencies of and increased variation in temperature and weather extremes, which can affect key biological processes, such as development, metabolism, and nervous system functioning and can lead to permanent shifts in range distribution (McCain and Garfinkel 2021) and phenology (Memmott et al. 2007; Vasiliev and Greenwood 2021; Rahimi and Jung 2024).

For insects, the timing of major life-stage events, such as development, growth, reproduction, and dormancy, is highly influenced by environmental conditions (M. J. Tauber and Tauber 1976). With respect to dormancy, strategies have evolved to allow taxa to avoid temporary periods of harsh climatic conditions with different stages of dormancy, including initiation, maintenance, and termination, tightly regulated by the endocrine system in response to environmental cues, such as photoperiod and temperature (D. L. Denlinger, Yocum, and Rinehart 2012). For example, in temperate climates, overwintering is a prevalent dormancy strategy and can occur during the adult stage, where metabolism is reduced or during developmental stages, where, additionally, development is halted (David L. Denlinger 2022). In most cases, overwintering represents a prolonged period of non-feeding, leaving dormant insects reliant on nutritional resources acquired before dormancy entry (Hahn and Denlinger 2007). Thus, in such states, overwintering insects take advantage of cooler ambient temperatures as observed during winter months in temperate climates to assist in metabolic suppression for prolonged periods of time when resources are unavailable and energetic reserves are limited (Gillooly et al. 2001). Many dormant insects coordinate their overwintering duration to correspond with the availability of resources and mates in their local environment and are reliant on environmental cues, such as increasing environmental temperatures, to initiate emergence (David L. Denlinger 2022). Given the dependence of insects on environmental temperatures for dormany maintenance and termination, it is hypothesised that during periods of dormancy, insects may be particularly vulnerable to shifts in environmental conditions, in particular temperature, due to changes in climate conditions (Bale and Hayward 2010; Freimuth et al. 2022; Fründ, Zieger, and Tscharntke 2013). As a consequence of ongoing climate warming, elevated overwintering temperatures may place additional pressure on already finite resources leading to changes in body condition, as evident by increased weight loss during dormancy (Hahn and Denlinger 2007). Such changes may potentially impact the ability of organisms to maintain dormancy, leading to reductions in survival, or limiting available resources for energetically-costly post-dormancy activities, such as foraging and mating, thus, contributing to predicted losses in fitness. Increases in winter temperatures can also lead to shifts in emergence timing, which may potentially have detrimental consequences due to intraspecific and interspecific phenological decouplings (Gordo and Sanz 2005; Schenk, Krauss, and Holzschuh 2018; Freimuth et al. 2022). This is particularly true for species where both sexes go through seasonal dormancy but differ in their timing of emergence as phenological shifts induced by climate warming might differentially affect the sexes, giving rise to potential temporal mismatches in gonadal development and/or mate availability (Zonneveld 1996) with negative consequences for the population as a whole. Similarly, for insect pollinators, which are reliant on the availability of floral resources, which in turn are reliant on insect-mediated pollination for reproduction, interspecific phenological mismatches are predicted to adversely affect fitness of both interacting parties, with detrimental consequences for ecosystem stability (Kudo and Ida 2013; Renner and Zohner 2018).

Concerns about phenological shifts have been explored in solitary bees (Bartomeus et al. 2011), which are an important group of ecological and commercial pollinators (Hoehn et al. 2008; Woodcock et al. 2019). Despite accounting for more than 85% of documented bee species (Danforth et al. 2019), solitary bees are generally understudied in contrast to social bees (T. J. Wood et al. 2020), where dormant states, such as diapause, have either been lost or reduced (e.g., honey bees) or are limited to one sex (e.g., bumblebees) (Santos, Arias, and Kapheim 2019; Colgan et al. 2019; Zapata-Hernández et al. 2024). Furthermore, as colony-level actions of social bees, such as nest thermoregulation, are predicted to help mitigate and reduce the damaging effects of environmental challenges (Straub et al. 2015), studies on thermal tolerance in such species may not be transferable to or representative of bees as a whole. Solitary bees may have increased susceptibility to environmental challenges given that, in many temperate species, both sexes must successfully undergo and complete overwintering before post-dormancy copulation occurs. Given that sexes differ in many aspects of their biology, including behaviour, physiology, morphology, and life-histories, differential responses to environmental stimuli may lead to phenological mismatches between the sexes. In particular, males may face further risks. Not only do many solitary bee species display protandry, meaning earlier emergence as a consequence of temperature exposure, may result in males emerging into less suitable environments. In addition, like other Hymenoptera, males are also haploid with sex determination being haplodiploidy with females developing from fertilised, diploid eggs and males developing from unfertilised, haploid eggs (Crozier 1977). The haploid nature of males means they automatically express alleles they carry (Joseph and Kirkpatrick 2004) meaning that males are potentially more prone to environmental pressures (O’Donnell and Beshers 2004) albeit evidence is still inconclusive (Ruiz-González and Brown 2006; Friedli et al. 2020; McAfee et al. 2022).

A common group of solitary bees that have been increasingly used to study the effects of environmental pressures during and after overwintering are mason bees (genus: *Osmia*) (J. Bosch and Blas 1994; Jordi Bosch and Kemp 2004; Fliszkiewicz et al. 2012; Schenk et al. 2018; Albacete et al. 2023). Mason bees are a taxon-rich group with approximately 350 described species, which are widely distributed in temperate climates (Banaszak and Romasenko 1998) and have gained increased attention due to their potential as efficient pollinators in agriculture, particularly for fruit crops (Gruber et al. 2011; Sedivy and Dorn 2014; Fliszkiewicz and Giejdasz 2023). Most mason bees are univoltine and undergo seasonal diapause in the summer as larvae, before spinning a cocoon, completing metamorphosis and entering a winter dormancy as adults, with emergence, mating, nest building and offspring provisioning taking place in the following spring when conditions become more favourable (Splitt, Schulz, and Skórka 2022). During winter dormancy, overwintering adults are solely reliant on resources provided to them prior to dormancy entry (J. Bosch and Kemp 2000). As most mason bees are above ground cavity nesters (Jordi Bosch, Maeta, and Rust 2001; Praz et al. 2008), temperature has a stronger influence on overwintering duration compared to other environmental factors, such as photoperiod (Beer et al. 2019). Elevated constant or monthly stepwise increases in temperature exposure can affect the duration of dormancy, weight loss and survival in overwintering mason bees, such as *Osmia bicornis*, *Osmia cornuta*, and *Osmia lignaria* (J. Bosch and Blas 1994; Jordi Bosch and Kemp 2004; Fliszkiewicz et al. 2012)(Schenk et al. 2018; Albacete et al. 2023). However, as above ground cavity-nesting bees, *Osmia* species are exposed to diurnal and seasonal fluctuations across their overwintering period. Therefore, there is a need to improve our understanding of how these organisms respond to temperature regimes informed by field-based recordings, which incorporate the variation in thermal exposures that are more reflective of what bees encounter during overwintering in their natural landscapes.

In addition, determining their reactions to thermal patterns as expected under current climate change scenarios is also urgently warranted (Paaijmans et al. 2013). Indeed, exposure to fluctuating temperatures during the prepupal larval stage has been shown to accelerate development in mason bees prior to diapause entry compared to constant temperature exposure (J. Bosch and Kemp 2000; Radmacher and Strohm 2011), highlighting the impact temperature regime can have on key ontogenetic events. In addition to development, elevated temperature is proposed to directly affect phenological shifts in terms of emergence timing of *Osmia* species yet our understanding of emergence patterns is limited as previous studies either artificially stimulated emergence through prolonged exposure at elevated temperatures (Kemp and Bosch 2005; Fründ, Zieger, and Tscharntke 2013) or used stepwise exposure regimes (Schenk et al. 2018). However, sudden stepwise increases of temperature can lead to mass emergence events, as during late-stage periods of dormancy, such as post-diapause quiescence, bees may be more sensitive to environmental temperature changes. In general, our understanding of the temperatures at which mason bees naturally emerge, whether it occurs at a critical thermal threshold in response to the cumulative effects of prolonged temperature exposure, or is connected with changes in intrinsic aspects of body condition, is still poorly understudied. In addition, information on how sexes, which naturally differ in phenology but are required to synchronise emergence to maximise individual fitness (Strobl et al. 2019), respond to current and future predicted overwintering temperature, is also still lacking. Similarly, many previous studies used bees from a single local source, which likely underestimates the phenotypic variability in thermal responses found in mason bees.

To address these limitations in our current understanding, we exposed mason bees to field-recorded temperature regimes, incorporating both diurnal and seasonal fluctuations, which were informed by projections for two potential climate change scenarios, which form the basis of the sixth and latest assessment report of the Intergovernmental Panel on Climate Change (Intergovernmental Panel on Climate Change (IPCC) 2021). According to these projections, surface air temperatures over land in Central Europe - relative to the 1995-2014 reference period - are projected to rise by around 3°C by 2100 under the intermediate-emission scenario SSP2-4.5 and by around 6°C under the worst case-emission scenario SSP5-8.5 (Intergovernmental Panel on Climate Change (IPCC) 2021; Tebaldi et al. 2021). Using an experimental approach, we investigated how temperatures based on such models affect dormancy emergence timing, condition (measured by relative weight loss) and survival in both sexes of two common mason bees, *O. bicornis* and *O. cornuta*, sourced from four geographically distinct locations per species. Given the accelerative effect that temperature exposure is predicted to have on overwintering bees in terms of emergence and conditional change, we predict that increased temperature exposure will correspond with earlier emergence, elevated weight loss, and lower survival rates (Fig. 1). Additionally, to understand the relationship between temperature exposure, weight loss, and emergence timing, we hypothesised that, if emergence is dependent on the depletion rate of nutritional reserves, shifts in emergence timing caused by temperature exposure would be strongly mediated by increased weight loss prior to emergence.

**Figure 1.**
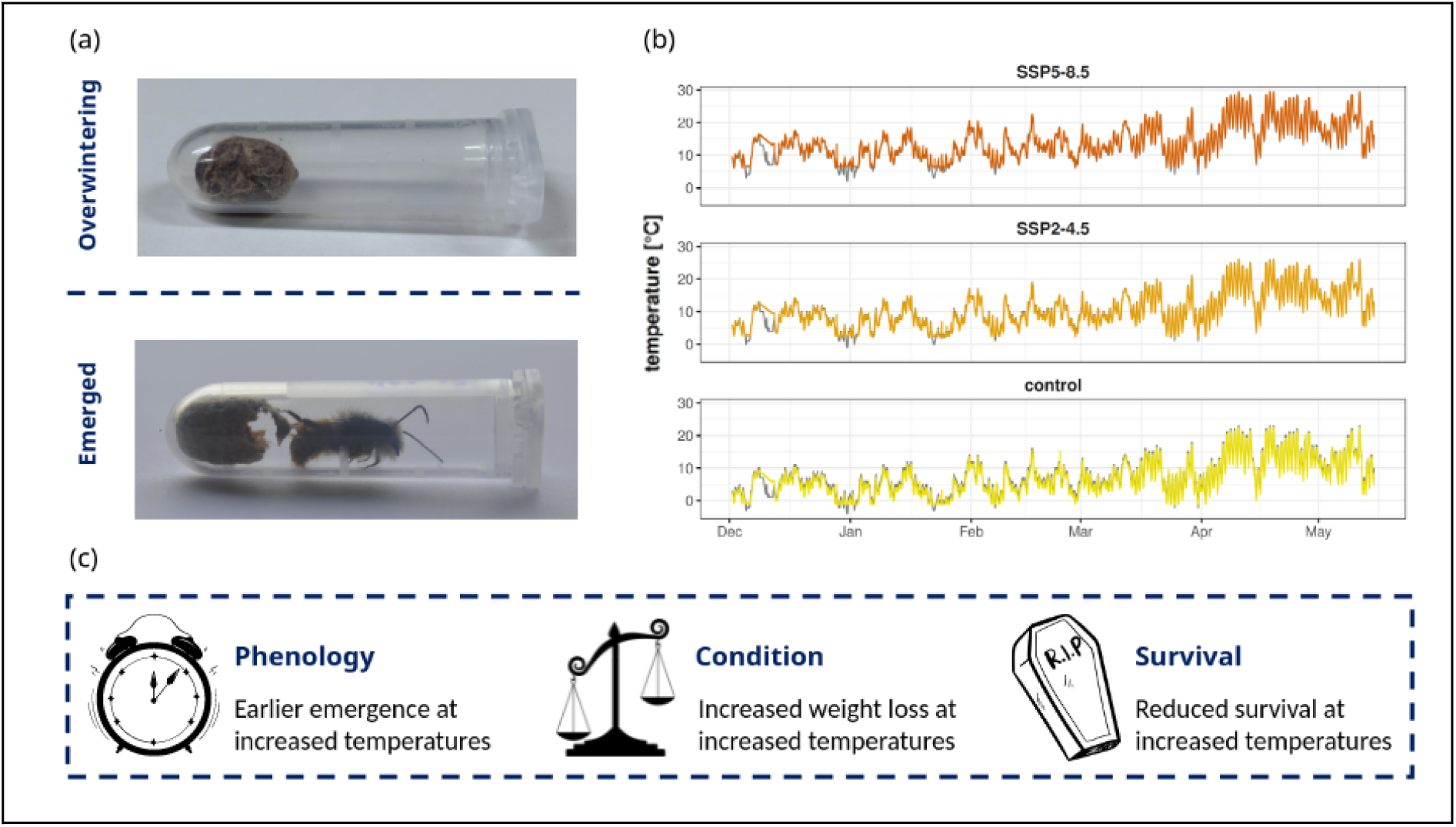
Experimental set-up and simulated temperature profiles based on current and predicted climate change models. (a) Images of individuals of the European orchard bee (*Osmia cornuta)* overwintering in a cocoon (upper image) or freshly emerged after overwintering (lower image). (b) Three line graphs depicting an overlay of the three temperature profiles simulated in our experiment (grey) that mason bees were exposed to and their achieved implementation in the form of logged temperatures for each temperature treatment (yellow = control, orange = SSP2-4.5, red = SSP5-8.5). The lower plot facet includes the control group (‘current’) temperature profile, representing surface air temperatures measured 2m above the ground at the weather station of the German National Meteorological Service in Mainz-Lerchenberg, Germany in the winter of 2019/2020. The upper two plots include the same profile, with each time-point increased by a fixed 3°C and 6°C relative to the control group profile, representing the climate change scenarios SSP2-4.5 (“intermediate scenario”) and SSP5-8.5 (“worst-case scenario”), respectively, that were laid out by the Intergovernmental Panel on Climate Change in their sixth assessment report. (c) Conceptual portrayal of the three main predictions that we made in relation to how bees respond to increased temperature scenarios.

## Materials and Methods

### Sampling and handling of individuals

We ordered 1153 mason bee cocoons, composed of cocoons of *Osmia bicornis* (n = 568) from four different commercial suppliers in Germany and Austria (Germany: Konstanz, Stahlberg, Greifswald; Austria: Schärding) and cocoons of *Osmia cornuta* (n = 585) from three different commercial suppliers (n = 445; Germany: Konstanz, Stahlberg; Austria: Schärding), complemented by cocoons of *O. cornuta* extracted from nesting boxes in Rottleberode, Germany (n = 140; approximate site coordinates are provided in Supplemental Table 1). All cocoons were received in October 2022. Upon reception, the cocoons were placed outside in a water-proof, outdoor cabin until the start of the experiment on December 1st, 2022. Before starting the experiment, cocoons were washed to remove nesting debris under running tap water and dried overnight in the laboratory at 25°C to avoid weight loss bias from debris sticking to the cocoon surface.

### Experimental setup

To simulate overwintering temperatures including predicted temperature increases under climate change, we used three programmable climate cabinets of the same model (RUMEDⓇ Type 3201, Germany). Each climate cabinet was assigned one of three temperature profiles. The temperature profiles consisted of a baseline profile representing current, field-relevant winter temperatures (as measured by the German National Meteorological Service (DWD) weather station in Mainz-Lerchenberg, Germany between December 2019 and June 2020) and two experimental treatment profiles based upon projections of surface air temperature increases over land in Central Europe by the year 2100 (Intergovernmental Panel on Climate Change (IPCC) 2021; Tebaldi et al. 2021) under two Shared Socioeconomic Pathway climate warming scenarios (intermediate emissions: SSP2-4.5, worst-case emissions: SSP5-8.5), which we approximated by implementing a fixed 3°C and 6°C temperature increase relative to the baseline temperature profile, respectively. Similarly to other studies (Ostap-Chec et al. 2021; Ruddle et al. 2018; Nicholls et al. 2017; Szentgyörgyi et al. 2017; Coudrain et al. 2016; Peters, Gao, and Zumkier 2016; Kehrberger and Holzschuh 2019), we used reference temperatures (and temperature projections), measured at 2m above ground, as both *O. bicornis* and *O. cornuta* nest above-ground in a variety of pre-existing nesting substrates (Danforth et al. 2019). We programmed the climate cabinets to adjust the temperature every 3 hours according to the underlying weather data at a resolution of 1°C. We independently measured temperature and humidity in all three climate cabinets using individual data loggers (TFA DOSTMANNⓇ LOG32TH, Germany) throughout the duration of the experiment. An overview of the simulated temperatures used in each treatment group, including the temperatures recorded by the data loggers, across the duration of the exposure experiment, is depicted in Figure 1 (recorded data are provided in Supplemental Table 2).

Before the temperature exposure experiment, we weighed a 1.5ml microtube for each sample, which contained ventilation holes in the lid. The weight of these tubes was assumed constant throughout the experiment. Subsequently, a cocoon was placed inside each tube and measurements were repeated. To automate the weighing process and make it more robust against errors, we set up a microbalance (PESCALEⓇ MYA 5, precision: 0.1µg) to automatically transfer measurements to a computer via a serial interface upon the push of a button.

After initial measurements were completed, samples (tubes containing cocoons) were assigned to the three climate cabinets using block randomisation, ensuring species, sexes and source sites were uniformly distributed across the three climate cabinets. Within each climate cabinet, samples were assigned onto 80-well microtube racks, again using block randomisation, ensuring a uniform distribution of species, sex, and source site sources across racks, thus preventing a positional bias inside climate cabinets (further details on the randomised assignment of individuals is provided in the Supplemental Methods S1). In addition, the positions of the microtube racks inside the climate cabinets were rotated weekly during weight measurements to further avoid potential positional biases.

### Weight measurements and emergence tracking

The experiment was started on December 1st with the initial measurements of sample weights, which served as the first time point of the experiment. As noted above, samples were not weighed directly, but instead cocoons and microtubes were weighed together with tube weight subtracted later during data analysis. For a subset of samples (n = 873, total = 1153), we measured weight weekly on the same day each week to track when divergences in weight occurred between treatment groups while for the rest of the samples, weight was measured only at the start and end of the experiment (after hatching).

Upon the first hatching, which was recorded in January 2023, all cocoons were checked once per day every day until the completion of the experiment. Upon hatching, which was observable by the complete emergence of the adult from their cocoon, we recorded the date of hatching and weighed each bee on their respective emergence day (each emerged bee was measured inside their respective microtube, which also included the remnants of their cocoon). Using both the starting weight and the weight at emergence, we calculated the total weight loss for each bee across the experiment. In addition to weight, we also recorded and confirmed sex and species based on morphological characteristics.

On May 15th (23 weeks into the experiment), as hatching had drastically decreased over the preceding weeks and with over 85% of bees emerged, the experiment was stopped and temperatures were raised to 26°C across all three climate cabinets for 24 hours to stimulate the emergence of remaining, alive individuals (J. Bosch and Blas 1994). Subsequently, all unhatched cocoons were opened via excision using microscissors. For each excised individual, we recorded species, sex, developmental stage (pupa/prepupa/imago), survival status at retrieval (alive/dead) and if applicable, parasite presence and genus. As expected from univoltine species, none of the unhatched mason bees were alive upon excision.

### Statistical analysis

Once all data were collected, we pre-screened the data and removed outliers if emergence weight was recorded to be higher than initial weight, which resulted in the removal of seven samples, or if a sample had a sudden increase in weekly measured weight, which resulted in the removal of another six samples. One sample was removed because the cocoon was empty upon excision and 19 samples were removed because they were parasites. Thus, in total, 33 samples were removed during pre-screening (2.86%, total = 1153), leaving a dataset of 1120 samples of which 273 were weighed only at the start and end of the experiment and 847 were additionally weighed weekly throughout the experiment.

In our analysis to investigate the effect of elevated temperature exposure on weight loss and emergence, we further reduced the pre-screened dataset by including only samples that represented individuals that had hatched alive before the end of the experiment (87.20%, n = 997, total = 1120). We fitted two linear mixed-effects models (R package lme4; (Bates et al. 2014)) to investigate the effect of temperature on emergence and weight loss, respectively. In the model that included total weight loss as the response variable, we defined weight loss as the difference in percentage of body weight at emergence relative to body weight at the start of the experiment to account for natural differences in body weight between individuals, sexes, and species. In both models, temperature treatment, species, and sex were included as interaction variables and source site was always included as a random effect. To test, when differences in weight loss became noticeable, we used the subset of 847 weekly-weighed samples and employed a Generative Additive Mixed-Effects Model (R package mgcv; (S. N. Wood 2010)) with weight loss as a dependent variable, temperature treatment, species, and sex as independent interaction variables and source site as a random effect.

To test differences in survival rate between temperature treatment groups, we used the full pre-screened dataset including both alive and dead individuals and employed a generalised linear mixed-effects model (R package lme4; (Bates et al. 2014)) to fit a binomial distribution with survival as the response variable, and temperature, species, and sex included as interaction variables and source site as a random effect.

We also tested whether weight loss as a mediator variable explained the effect of increased temperature treatments on emergence timing. To this end, we implemented a moderated mediation analysis (R package mediate; (Yimer et al. 2023)) that compares a full model that includes the proposed mediator variable against a reduced model that lacks the mediator variable (Hayes 2017). The full model contained temperature treatment as an independent variable, emergence timing as a dependent variable and weight loss as a mediator. Analogously to the linear models, all experimental group combinations of species and sex were additionally included as interaction effects, which in this context are also referred to as moderator variables (Preacher, Rucker, and Hayes 2007).

Lastly, to test the hypothesis that heavier or larger individuals emerged before lighter individuals, we again implemented two linear mixed effects models identical to the first two models, but additionally including either initial body weight or body length as predictor variables. Like for the first two models for emergence timing and weight loss, the pre-screened dataset was used, but individuals that had not hatched alive were excluded.

For all models, differences within and across experimental groups were assessed using Tukey-Kramer post-hoc tests on estimated marginal means (R package emmeans; (Searle, Speed, and Milliken 1980)) and corrected for multiple testing using the Tukey adjustment method and the statistical significance of interaction effects was evaluated using Type-II Wald Χ² tests (R package car; (Fox and Weisberg 2018)) and corrected for multiple testing using the Bonferroni-Holm’s adjustment method.

For verification, validation, and reuse, all steps of the statistical analysis were annotated and are provided in Supplemental Script 2 and raw data for the statistical analysis are provided in Supplemental Table 3.

## Results

### Early emergence under elevated temperatures

To determine the effects of temperature on shifts in emergence timing, we implemented three field-based overwintering temperature regimes that included seasonal and diurnal variation, to which we exposed females and males of both *Osmia bicornis* and *Osmia cornuta*. We observed significantly earlier emergence under both elevated-temperature scenarios, a pattern which was consistent across all combinations of species and sex (Linear Mixed Effect Model (LMM); Tukey-Kramer test: Tukey-adjusted P < 0.05; Fig. 2). Across all bees, mean difference (MD) in date of emergence was 40.54 days earlier in the worst-case emission group and 18.84 days earlier in the intermediate emission group, with standard errors (SE) of 1.21 days and 1.20 days, respectively. An interaction effect of temperature, species and sex was significant (LMM; Type-II Wald Χ² test; Bonferroni-Holm-adjusted P < 7.8e-10), prompting us to explore species and sex differences in more detail. As expected, due to known protandry in these systems, in the control group and in both elevated temperature groups, males, on average, emerged more than one week earlier than females (LMM; Tukey-Kramer test: Tukey-adjusted P < 0.05). However, in *O. bicornis* the difference in emergence timing between the sexes was increased by an additional 5.91 and 8.18 days in both treatment groups, respectively, compared to the control group (control: 9.68 (MD) ± 0.74 days (SE); SSP2-4.5: 15.59 (MD) ± 1.07 days (SE); SSP5-8.5: 17.86 (MD) ± 1.43 days (SE)). In contrast, for *O. cornuta*, we did not observe this effect as sex differences were inconsistent across treatment groups with respective to timing of emergence (control: 10.58 (MD) ± 1.09 days (SE); SSP2-4.5: 12.81 (MD) ± 1.17 days (SE); SSP5-8.5: 8.30 (MD) ± 1.09 days (SE); Fig. 2). In relation to species differences, *O. cornuta* individuals, on average, emerged 3.5 weeks before *O. bicornis* (females: 27.42 (MD) ± 1.63 days (SE); males: 24.48 (MD) ± 1.60 days (SE)). In addition, we observed that for *O. bicornis,* individuals emerged on days around the same mean temperature (control: 16.4 ± 0.1°C (SE); SSP2-4.5: 16.3 ± 0.2°C (SE); SSP5-8.5: 16.0 ± 0.2°C (SE)) with no significant differences between treatments (LMM; Tukey-Kramer test: Tukey-adjusted P > 0.05), while for *O. cornuta* mean temperature at day of emergence (control: 11.0 ± 0.2°C (SE); SSP2-4.5: 11.7 ± 0.2°C; SSP5-8.5: 14.7 ± 0.2°C) varied significantly with temperature treatment (LMM; Tukey-Kramer test: Tukey-adjusted P < 0.05).

**Figure 2.**
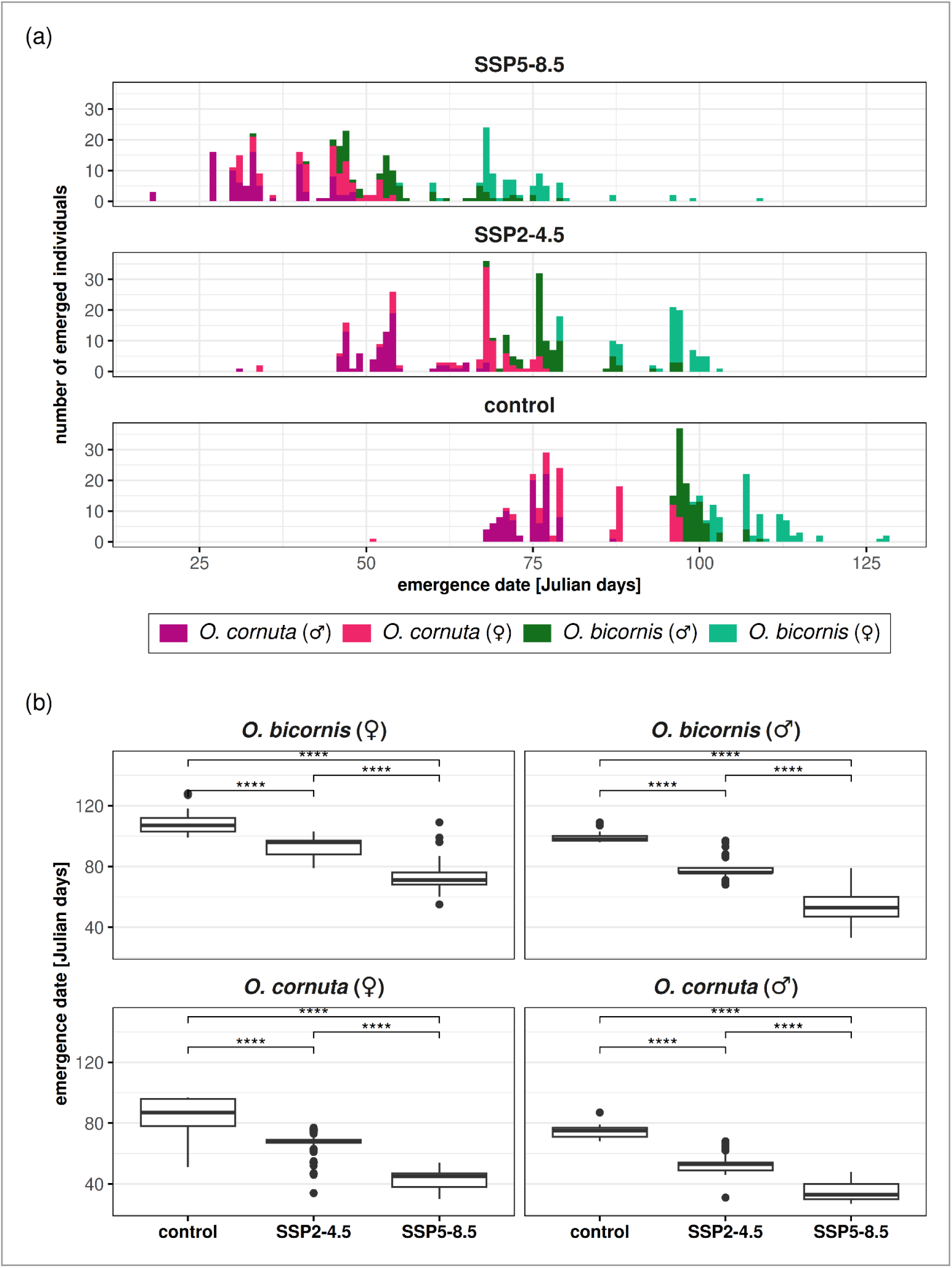
Earlier emergence with increasing overwintering temperature exposure. (a) Stacked bar plots showing for all species and sexes the number of daily emerged individuals (y-axis) in relation to the time of emergence (x-axis), as measured in Julian days since 1st of January. The lower plot facet depicts the emergence patterns observed with the control group temperature profile, while the upper two facets show emergence patterns under climate change scenarios SSP2-4.5 and SSP5-8.5, respectively. Each species and sex are designated by an individual colour. (b) Box plots representing distributions of emergence times (y-axis) for each species and sex (facets) within each temperature treatment (x-axis). For each species and sex, pairwise differences between temperature treatments are shown with level of significance indicated by number of asterisks (LMM, pairwise contrasts of estimated marginal means using Tukey-Kramer test, adjusted for multiple testing: ns = not significant, * = P < 0.05, ** = P < 0.01, *** = P < 0.001, **** = P < 0.0001).

### Survival rates during overwintering were not generally affected by elevated temperatures

While earlier emergence and weight loss may have indirect effects on fitness, we also recorded survival rates for all bees in our experiment. Using all bees, we found no significant difference in survival rates across temperature treatments (Generalised Linear Mixed Effect Model; Tukey-Kramer test: Tukey-adjusted P > 0.05) highlighting the ability of both species to at least survive and emerge from prolonged elevated thermal exposure. Despite this, irrespective of temperature exposure, we found that *O. bicornis* bees generally had a significantly lower survival rate (87.3%) than *O. cornuta* (91.3%, Fisher’s Exact Test, P = 0.033, Fig. 3). Furthermore, within *O. bicornis*, males (84.3%) had a significantly lower survival rate compared to females (90.8%, Fisher’s Exact Test, P = 0.028). In *O. cornuta*, we also observed a lower survival rate in males but this result was not statistically significant (males: 89.2%; females: 93.8%; Fisher’s Exact Test, P = 0.071).

**Figure 3.**
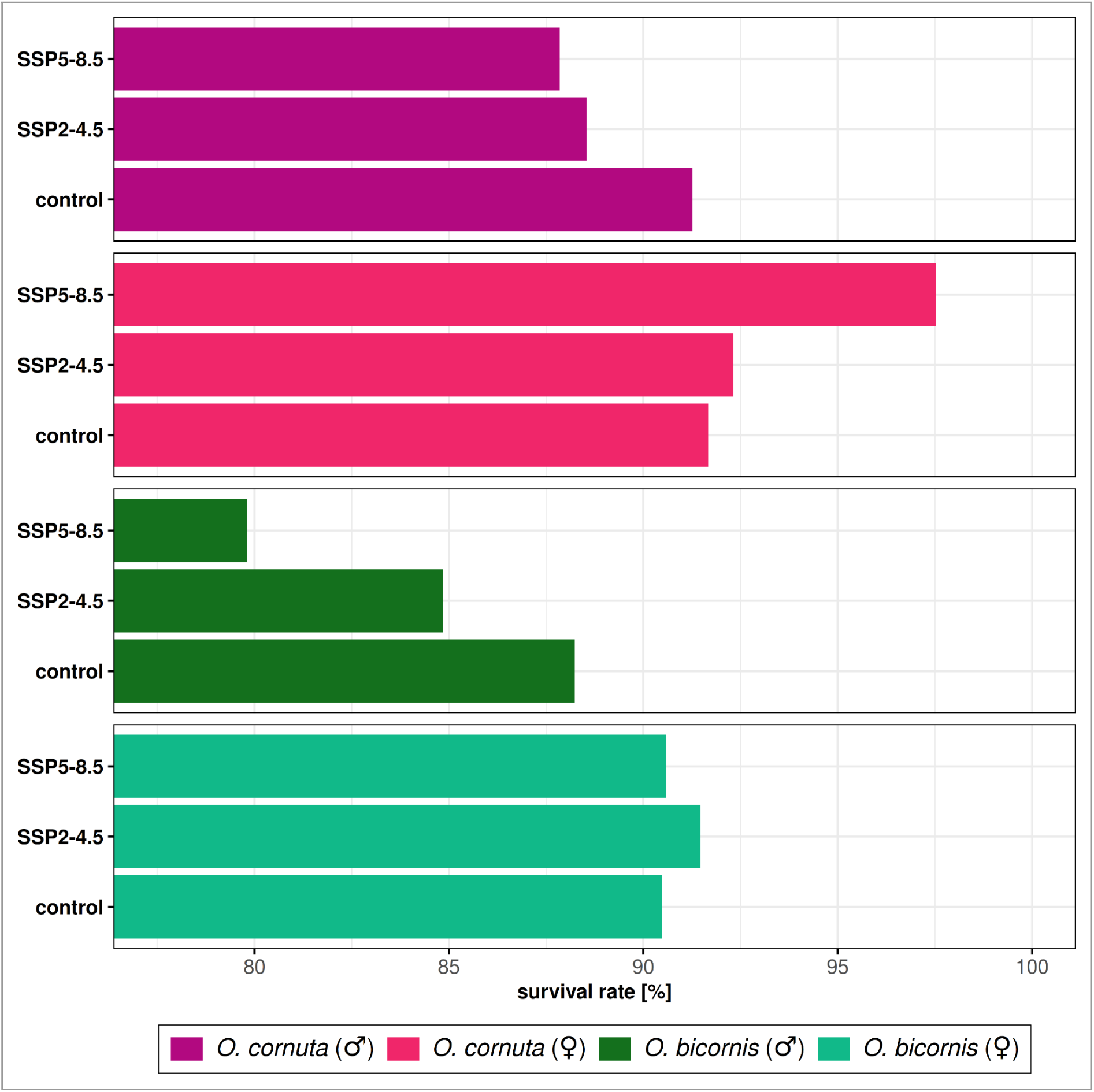
Differences in survival in response to elevated overwintering temperatures. Bar plot showing survival rates (x-axis) across all temperature treatments (y-axis) for each species and sex. Survival rates across temperature treatment groups were not significantly affected by elevated temperatures (Generalised Linear Mixed Effect Model: Bonferroni-Holm adjusted P = 0.53), however, survival rates in *O. bicornis* (87.3%) were significantly lower than in *O. cornuta* (91.3%, Fisher’s Exact Test, P = 0.033) and were also significantly lower in *O. bicornis* males (84.3%) compared to O. *bicornis* females (90.8%, Fisher’s Exact Test, P = 0.028).

### Increased weight loss with elevated temperature exposure

As weight loss due to a reduction in nutritional reserves has been posited as a factor associated with earlier emergence, we also investigated the effect of temperature exposure on weight loss. We observed a significant increase in weight loss for both elevated temperature exposure groups across all species and sex combinations (LMM; Tukey-Kramer test: Tukey-adjusted P < 0.05, Fig. 4) with the strongest effect observed in the worst-case emission group where mean weight loss at emergence was 1.49 times higher compared to the control group (control: 7.76 ± 0.15% (SE); SSP5-8.5: 11.60 ± 0.19% (SE)). In comparison, increased weight loss was also observed in the intermediate emission group but at a more moderate level (1.17 times higher) compared to the control group (SPP2-4.5: 9.09 ± 0.19% (SE)). In addition to comparing total relative weight loss, during the experiment, we also tracked changes in weight for a subset of bees (75.72%, n = 873 individuals) each week to determine when changes in weight loss occur. We found increasing differences in weight loss across time when comparing elevated temperature groups to the control group, which were significant after two weeks of weight tracking in the worst-case emission group and after ten weeks in the intermediate emission group (Generative Additive Mixed Effect Model; overlap in 95% CIs) highlighting that not only did exposure to elevated temperatures result in elevated weight loss but the observed timing of the effect was considerably earlier in the worst-case scenario group.

**Figure 4.**
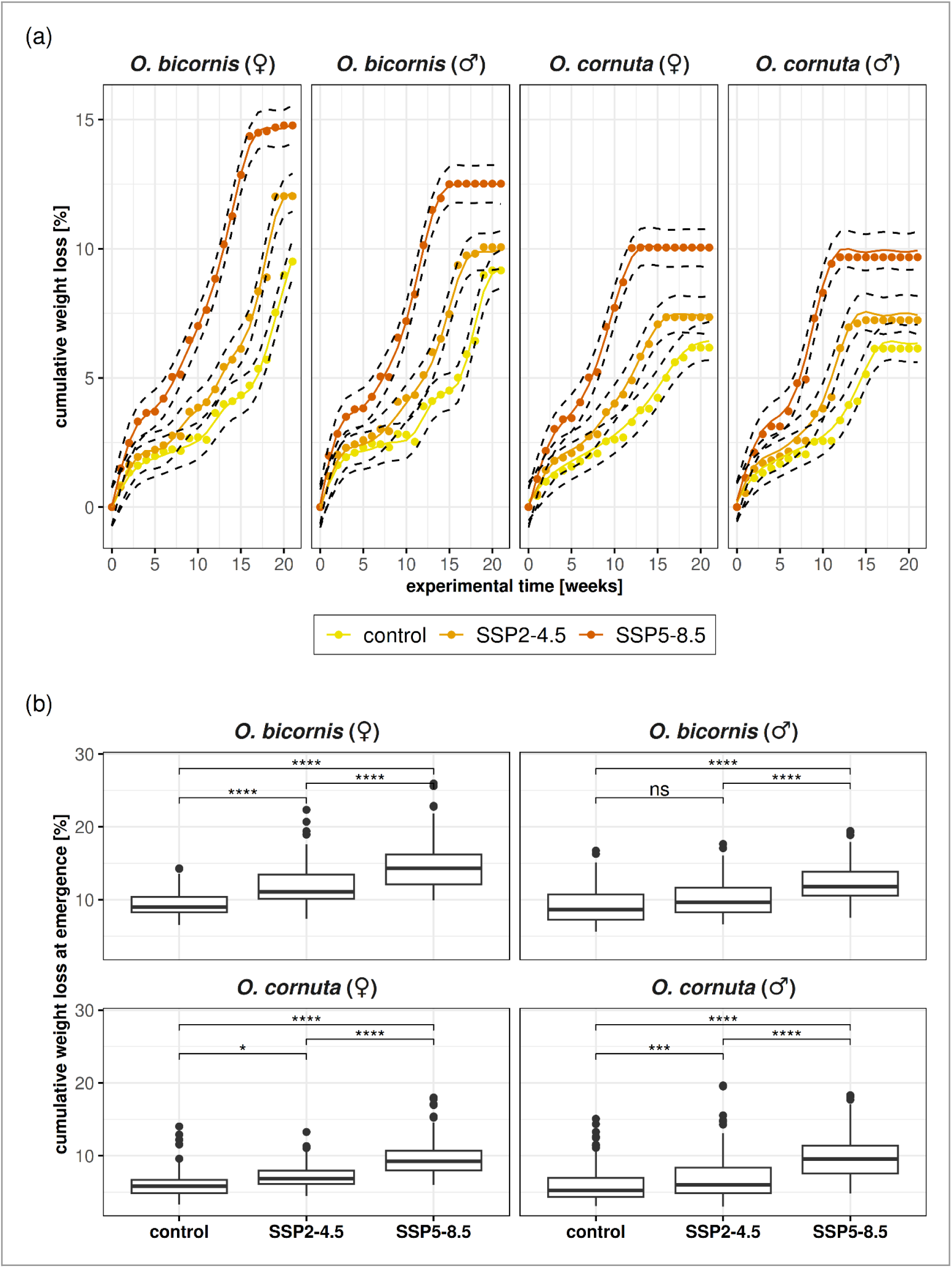
Increased weight loss in response to elevated overwintering temperatures. (a) Line plots depicting the accumulation of weight loss (x-axis) over experimental time in weeks (y-axis) during the overwintering experiment for each species and sex (facets) and temperature treatments (yellow = current, orange = SSP2-4.5, red = SSP5-8.5). Coloured dots represent weekly mean weight loss, averaged across individuals within the same group, while coloured lines represent approximations of that raw data fitted using generative additive mixed effect models with dotted black lines marking the upper and lower bounds of the 95% confidence intervals of the model predictions. (b) Box plots representing distributions of cumulative weight loss at emergence (y-axis) for all temperature treatments (x-axis), including species and sexes (facets). For each species and sex, pairwise differences between temperature treatments are shown with level of significance indicated by number of asterisks (LMM, pairwise contrasts of estimated marginal means using Tukey-Kramer test, adjusted for multiple testing: ns = not significant, * = P < 0.05, ** = P < 0.01, *** = P < 0.001, **** = P < 0.0001).

### Effect of temperature on emergence timing is only partially mediated by weight loss

As the role of weight loss has been suggested as a potential contributor to emergence in overwintering insects, we examined if, and by how much, early emergence in response to temperature exposure was mediated by weight loss. We found a significant mediation effect for weight loss at emergence (Moderated Mediation Analysis, Average Mediated Effect = 9.90, Average Direct Effect = 51.33, P < 10⁻¹⁶), however, the proportion of the effect of temperature exposure on emergence timing mediated by weight loss amounted to only 23.9%, indicating that the effect of temperature on emergence timing is mediated by weight loss only to a limited extend.

### Size and body weight are only weak predictors of emergence timing

It has also been suggested in the literature that larger individuals emerge significantly earlier than smaller individuals in both *O. cornuta* and *O. bicornis* as earlier emergence might be beneficial in terms of avoiding competition over resources. We, thus, investigated the relationship between body size, initial body weight before the start of the experiment and weight loss, as well as emergence timing, to understand if differences in response to temperature exposure are predicted by body weight or size. Contrary to what has been reported prior, we found that in both species heavier or larger individuals tended to emerge later than lighter or smaller individuals in response to increased temperatures (Supplemental Fig. 1). Overall, we found that initial body weight and body size were significantly associated with both increased weight loss at emergence and earlier timing of emergence (LMM; Type-II Wald Χ² tests: Bonferroni-Holm-adjusted P < 0.05), however, such findings were not consistently significant across species, sex, and treatment combinations (LMM; Tukey-Kramer test: Tukey-adjusted P < 0.05). In addition, the contribution of initial body weight in explaining variation in emergence measured in terms of change in coefficient of determination (ΔR²) was extremely low as removing body weight from the model reduced the variation explained by only 0.1% (full model: R² = 94.72%; reduced model: R² = 94.62%). The same effect was observed when exchanging initial body weight for body size. Together, this indicates that even if body condition contributes to the variation in emergence timing across and within experimental groups, the effect of its contribution is likely small or negligible.

## Discussion

In the face of ongoing climatic change, understanding the responses of organisms to increasing elevated temperatures represents one of the most pressing and timely challenges. While advancements are being made in understanding thermal tolerance, we urgently need to assess the capacities of ecologically essential species, such as wild bees, to cope with current field-relevant temperatures and temperatures predicted under future climate scenarios. We, thus, exposed overwintering mason bees to temperature regimes informed by recent historical records and models developed by the Intergovernmental Panel on Climate Change to determine how elevated temperature exposure during dormancy affects timing of emergence, survival, as well as weight loss. We found that exposure to temperatures predicted under intermediate and worst-case scenarios associated with climate change had strong effects on phenology with both species emerging earlier with increasing elevated temperature exposure. Elevated temperature also led to increased divergence in emergence timing between male and female *O. bicornis*, indicating potential intraspecific phenological mismatches. In addition, bees also lost increasingly more weight at higher temperatures yet despite these losses, both weight loss and body size, were weak predictors of premature emergence. While elevated weight losses may have consequences for post-dormancy survival and fitness, we found no decrease in survival rates across temperature regimes highlighting the capacity of mason bees to at least survive overwintering at prolonged temperature exposure.

For many overwintering species, the entry into and maintenance of dormant states allows for avoiding harsh environmental conditions where unfavourable climate may potentially limit mating opportunities (David L. Denlinger 2002) and food availability (Wilsterman, Ballinger, and Williams 2021; David L. Denlinger 2002). For overwintering insects, emergence timing is generally influenced by environmental factors, such as photoperiod (Adkisson 1964) and temperature (Maurice J. Tauber, Tauber, and Masaki 1986), meaning that shifts in seasonality as predicted under climate change may lead to shifts in overwintering termination with organisms emerging into potentially more inhospitable environment conditions that they are ill-prepared for (Van Dyck et al. 2015). Here, we found that the timing of emergence for both *O. cornuta* and *O. bicornis* was affected by increased temperature exposure. In particular, the strength of the effect of temperature was proportional to the temperature regime with bees exposed to temperatures predicted under scenarios SSP2-4.5 and SSP5-8.5 emerging nearly three and six weeks earlier, respectively. Previous studies on mason bees also found that elevated temperature exposure during overwintering led to earlier emergence from winter dormancy yet such studies either used constant (Kemp and Bosch 2005) or step-wise monthly temperature increases (Schenk et al. 2018), which lack the thermal variation that overwintering bees encounter in their natural environment, and which is predicted to increase the sensitivity of insects to climate change (Paaijmans et al. 2013). Similarly, a common procedure used in research on solitary bees involves the artificial stimulation of emergence through incubation of overwintering bees at elevated temperatures (e.g., 17-26°C (J. Bosch and Blas 1994; Jordi Bosch and Kemp 2003, 2004; Kemp and Bosch 2005; Fründ, Zieger, and Tscharntke 2013; Albacete et al. 2023)), which may possibly contribute to or exacerbate detrimental effects on post-dormancy performance. While such stimulation temperatures allow for regulation of the timing of emergence for the timed use of mason bees in agricultural practices (e.g., pollination in orchards (J. Bosch and Blas 1994)), it limits our understanding of phenotypic variation in terms of emergence, which is required when determining potential ecological mismatches at the intra- and interspecific levels, due to climate-mediated phenological shifts. Indeed, in our experiment, we found that bees routinely emerged at temperatures on average lower than artificial termination temperatures routinely used in the literature, highlighting that lower temperatures may be more appropriate as a stimulant for initiation of post-dormancy activities although further research is required to determine consequences that might exist.

While earlier emergence may be compensated for by the polylectic nature of mason bees (Bartomeus et al. 2011), the biodiversity-associated buffering capacity of an ecosystem (Bartomeus et al. 2013), complementary phenological shifts in floral resources (Rafferty and Ives 2011), or coordinated timing of mate emergence (Zonneveld 1996), the predicted lower diversity of plants under climate change scenarios (Harrison 2020) may still dampen mitigative actions. For example, earlier emerging bees may compensate by choosing a less optimal or new floral resource, which may have implications for long-term survival and fitness. Indeed, a previous semifield-based study found that even a three day phenological mismatch between mason bee emergence and floral blooming resulted in fitness losses (Schenk, Krauss, and Holzschuh 2018) underlining the negative connotations earlier emergence may have on mason bees. The concerns over potential interspecific mismatches are further highlighted by observations of earlier flowering in response to climate change-induced temperature increases (Menzel 2003; Gordo and Sanz 2010; Amano et al. 2010; Renner, Wesche, and Zohner 2021). Predictions from German-based studies (Renner, Wesche, and Zohner 2021), which are likely more relevant to our findings given our use of bees originating from sites in Germany and Austria, translate to earlier flowering by 9.6-12.6 days and 19.2-25.2 days under the SSP2-4.5 and SSP5-8.5 scenarios, respectively. Together with our findings of earlier emergence by a mean 19 days under the SSP2-4.5 scenario and a mean 41 days under the SSP5-8.5 scenario, this suggests that mismatches may be potentially greater than previously assumed (Bartomeus et al. 2011; Thackeray et al. 2016). Although a formal examination is required, the resulting mismatches of about one and two weeks under the intermediate and worst-case emission scenarios, respectively, would exceed mismatches that were previously found to be detrimental in terms of fitness for mason bees (Schenk, Krauss, and Holzschuh 2018), adding to concerns about phenological mismatches in response to climate change.

In addition to differences in emergence timing across different temperature exposure regimes, within *O. bicornis*, we found evidence of the sexes responding differently to elevated temperatures. More specifically, the difference in emergence timing between males and females increased significantly from 10 days in the control group to 16 and 18 days in the intermediate and worst-case emission scenarios, respectively. Previous reports for sex differences in emergence timing vary between four and 11 days for mason bees (J. Bosch and Blas 1994; Raw 2009) but with considerable variation depending on population or ecotype examined (Tasei and Picart 1973). The monandrous nature of mason bees where females are generally single-mated (Paxton 2005), places additional pressure on males to emerge earlier, forage, and locate a mate as in such systems, mating with multiple females is a strategy for increasing fitness (Seidelmann 2014). Mason bees also have complex behavioural repertoires during copulation meaning males must invest time and energy in differentiating receptive and non-receptive females (Strobl et al. 2019), meaning earlier male emergence could be advantageous. However, as emergence timing is predicted to be tightly linked between the sexes, intraspecific divergences, such as those observed here, may lead to predicted elevated sexual antagonism, which may result in fitness costs (Ekrem and Kokko 2023). Similarly, male emergence has been more strongly linked with flowering patterns (J. Bosch and Blas 1994) meaning that in situations where phenological mismatches occur between plants and pollinators, males may be disproportionately affected. Our findings indicate that under worst-case scenarios, intraspecific divergence in phenology may occur, which may have consequences at the population size. While such differences were not identified in *O. cornuta*, the species-specific responses may be a consequence of *O. bicornis* naturally emerging later from overwintering, and, therefore, being subjected to higher temperature exposure for a more prolonged period. While later emergence is also found naturally in the wild, it may indicate that later-emerging species may face more problems due to higher temperature exposure during overwintering.

Exposure to elevated temperature before and during winter dormancy has been associated with fitness losses, including increased mortality (Radmacher and Strohm 2011), reduced longevity post-emergence, and increased pesticide sensitivity (Albacete et al. 2023). Therefore, under the predicted climate change scenarios, lower survival rates would be predicted with increasing temperature exposure. Contrary to this prediction, we found no evidence that increased temperature exposure leads to higher mortality during overwintering, a finding in line with a previous study that exposed mason bees to different constant overwintering temperatures and found that survival was unaffected (Fründ, Zieger, and Tscharntke 2013). However, post-dormancy longevity was reduced in *O. cornuta* exposed to elevated overwintering temperatures that overlap with our intermediate scenario (Albacete et al. 2023), highlighting that non-lethal observations reported in our study may still have long-term consequences post-emergence in terms of longevity and associated fitness. Thus, while our finding indicates the capacity of overwintering mason bees to survive prolonged exposure at elevated temperatures, further research is required to determine if such exposure results in long-term fitness consequences after emergence.

Our study highlights the strong influence elevated temperature exposure can have on timing of emergence yet the exact triggers leading to dormancy emergence are likely to be a combination of extrinsic and intrinsic factors. Abiotic factors, such as temperature and photoperiod, affect the timing of winter dormancy emergence, yet, given the cavity-nesting nature of overwintering mason bees, the influence of photoperiod on emergence is considered to be less strong compared to temperature (Beer et al. 2019). In comparison, how intrinsic factors, such as changes in body weight, contribute to timing of emergence are less well understood. Here, we found exposure to elevated temperatures predicted under climate change scenarios SSP2-4.5 and SSP5-8.5 resulted in considerable increases in weight loss, with measurably losses detectable after just two weeks of exposure in bees designated to the worst-case scenario group. Although our measurements are based on whole-bodied individuals, including their cocoons, previous studies on mason bees have demonstrated that exposure to elevated temperatures leads to loss of fat reserves (Fliszkiewicz et al. 2012), while during overwintering, essential compounds, such as carbohydrates and lipids decrease (Dmochowska et al. 2013; Wasielewski et al. 2013) indicating that increased metabolic activity in response to elevated temperature may contribute to the weight loss described in our present study. Given the importance of such reserves to an individual’s post-dormancy survival and fitness, it is highly probable that elevated weight loss is negatively associated with fitness as has been described in previous studies that employed constant temperature exposures (Jordi Bosch and Kemp 2004).

As reductions in energetic resources may contribute to premature emergence (Short and Hahn 2023), we examined the relationship between temperature exposure, weight loss, and early emergence using a mediation model and found that, contrary to what we had hypothesised, weight loss was only a weak mediator of the effect of temperature exposure on emergence timing. Although this analysis does not replace the need for experimental studies to definitively assess this relationship, our results suggest that weight loss is unlikely to be the main physiological process driving emergence. While other environmental factors, such as changes in photoperiod, are associated with emergence in other insects from overwintering, the influence is predicted to be less in mason bees given they overwinter in cocoons within cavities (Wasielewski et al. 2013), and is instead more influential on post-emergence activities (Beer et al. 2019). In contrast, ontogenetic factors and pre-wintering temperatures (Jordi Bosch, Sgolastra, and Kemp 2010), likely contribute more to phenotypic variation observed within our analysis. Similarly, differences in phenotypic variation may be explained by genetic differences within and across source sites as has been found in other systems (Pruisscher et al. 2018) and represents a future avenue of research to understand the genetic bases of phenological variation across natural populations.

Aside from environmental factors and weight loss, it has been suggested that body condition, including initial body weight and body size, is an important predictor of variation in emergence timing and weight loss across individuals with heavier individuals suggested to emerge before lighter individuals in response to temperature exposure (Schenk et al. 2018). Earlier emergence in larger individuals has been hypothesised as a strategy to avoid competition over resources with heavier individuals possessing greater energy reserves needed to emerge earlier but also survive in potentially colder environments. We formally tested this hypothesis, finding that initial body weight was both associated with emergence timing, as well as weight loss, yet contrary to what has been reported prior, we found a pattern suggestive of lighter individuals emerging before heavier individuals and with lower relative weight loss. Overall, the effect size of body condition as a predictor of emergence timing was very weak. In line with this, an earlier study that examined the effect of temperature exposure on overwintering *O. cornuta* also included body weight as a predictor variable in their analysis (Jordi Bosch and Kemp 2004) but did not find a significant effect on emergence timing and significant effects on survival and longevity only in males, suggesting that body condition is unlikely to play a major role in explaining emergence timing differences.

## Conclusions

With increasing environmental temperatures predicted under anthropogenic-mediated climate change, understanding the responses of organisms to changing environmental conditions is urgently required. Here, using field-informed temperature regimes, we demonstrate how elevated temperature exposure, reflective of intermediate and worst-case climate change scenarios, significantly influences the phenology and condition of two ecologically and commercially important solitary bees. Our study also underscores the complexity of predicted climatic change-associated impacts on solitary bee species, highlighting intra- and interspecific variation in terms of thermal responses within species that share similar nesting ecology. Information on solitary bee thermal responses, in general, will greatly benefit from examination of more species that differ in terms of nesting ecology, developmental stage of dormancy, seasonal timing of dormancy, as well as duration of dormancy. In addition, further insight into the responses of contemporary populations will be gained through the application of field or semi-field experiments that can help to provide more ecologically relevant information on thermal responses (CaraDonna, Cunningham, and Iler 2018). Similarly, further understanding changes and responses at the physiological and molecular levels will provide improved insights into the phenotypic plasticity underlying thermal responses. Collectively, our findings contribute to the growing literature highlighting the impact that elevated temperature will have on the timing of key life-stage events, which may have implications at the individual level through emergence timing and increased weight loss, at the population level through intraspecific phenological mismatches between the coordinated emergence timing of sexes (Maurice J. Tauber and Tauber 1978) and potentially consequences at the community level through interspecific mismatches between pollinators and floral resources (Visser and Both 2005). Through understanding such processes, mitigation strategies can be developed and implemented to ensure the conservation of essential pollinator species and the maintenance of essential ecosystem services.

## Data Availability Statement

Data and scripts required for reanalysis will be provided as supplemental files and will be made publicly available on GitHub upon publication.

## Acknowledgments

We would like to thank the German Federal Environmental Foundation (Deutsche Bundesstiftung Umwelt, DBU) for supporting JSM with a doctoral scholarship (20021/743). In addition, we would like to thank the Chinese Scholarship Council (202206170038) for supporting the doctoral scholarship of HX. We also thank the support of the Eva Crane Trust (ECTA_20210915), which provided financial support for this project, and Dr. Eckart Stolle for the provision of nest boxes.

## Author Contributions

JSM, LL and TJC conceived the ideas and designed methodology; JSM, LL, DH, IK, HX and TJC collected the data; JSM analysed the data; JSM and TJC led the writing of the manuscript. All authors contributed critically to the drafts and gave final approval for publication.

## Supplemental Figures

**Supplemental Figure 1.**
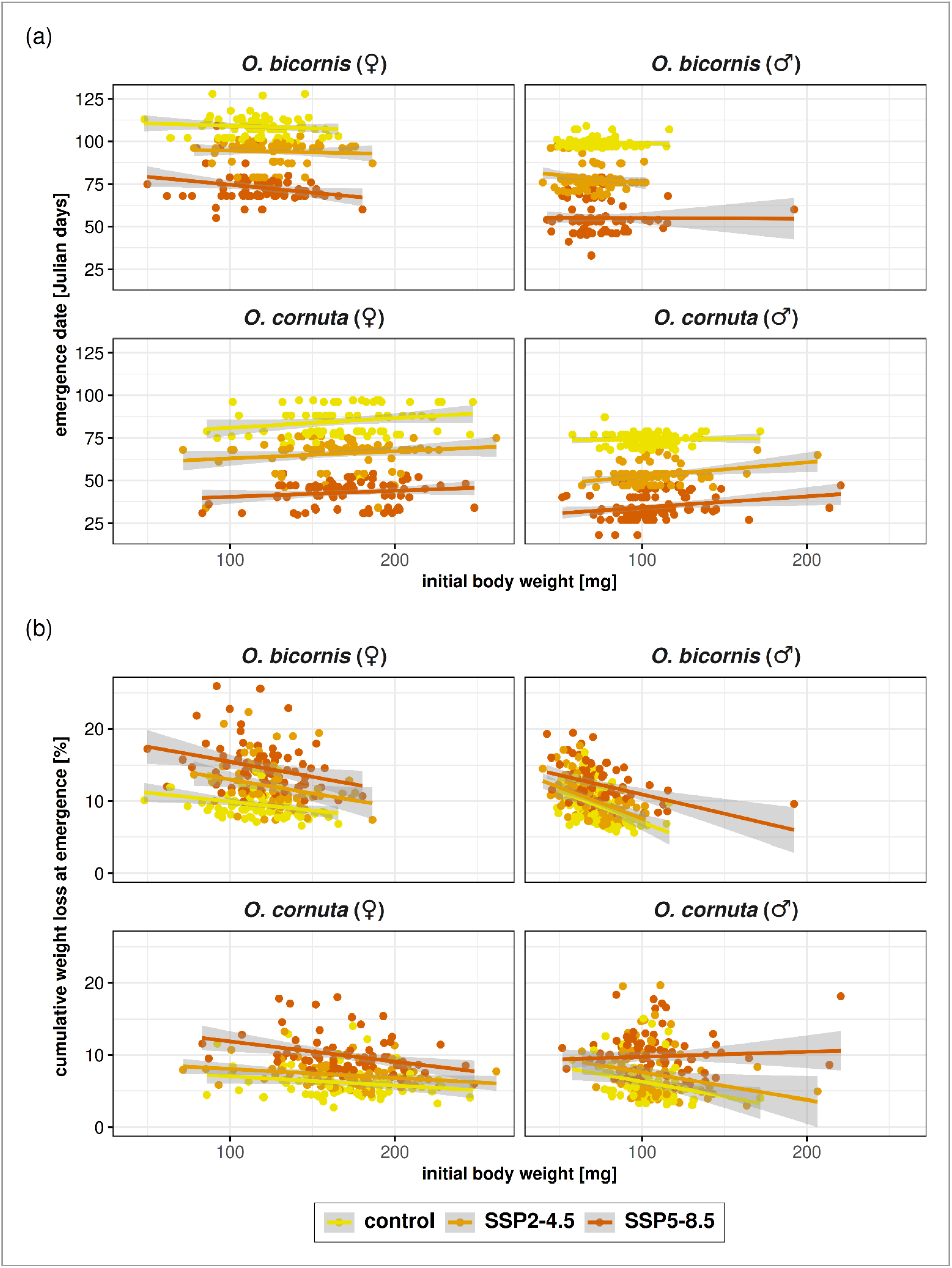
Effect of initial body weight on emergence timing and weight loss. Scatter plots depicting the effect of initial body weight (x-axis) on either: (a) emergence date; or (b) weight loss at emergence (both on y-axes) for all three treatment groups and species and sexes (facets). Colour-coded goodness-of-fit lines, representing each temperature group, are provided, with standard error of predictions in the background shown in grey.

